# Generation of heritable germline mutations in the jewel wasp *Nasonia vitripennis* using CRISPR/Cas9

**DOI:** 10.1101/096578

**Authors:** Ming Li, Lauren Yun Cook Au, Deema Douglah, Abigail Chong, Bradley J. White, Patrick M. Ferree, Omar S. Akbari

## Abstract

The revolutionary RNA-guided endonuclease CRISPR/Cas9 system has proven to be a powerful tool for gene editing in a plethora of organisms. Here, utilizing this system we developed an efficient protocol for the generation of heritable germline mutations in the parasitoid jewel wasp, *Nasonia vitripennis*, a rising insect model organism for the study of evolution, development of axis pattern formation, venom production, haplo-diploid sex determination, and host–symbiont interactions. To establish CRISPR-directed gene editing in *N. vitripennis,* we targeted a conserved eye pigmentation gene *cinnabar*, generating several independent heritable germline mutations in this gene. Briefly, to generate these mutants, we developed a protocol to efficiently collect *N. vitripennis* eggs from a parasitized flesh fly pupa, *Sarcophaga bullata*, inject these eggs with Cas9/guide RNA mixtures, and transfer injected eggs back into the host to continue development. We also describe a flow for screening mutants and establishing stable mutant strains through genetic crosses. Overall, our results demonstrate that the CRISPR/Cas9 system is a powerful tool for genome manipulation in *N. vitripennis*, with strong potential for expansion to target critical genes, thus allowing for the investigation of a number of important biological phenomena in this organism.

## Introduction

Hymenopteran insects, including all ants, bees, and wasps, represent one of the most prominent insect orders, occupying roughly 8% of all described species on earth^1^. The parasitoid wasp *Nasonia vitripennis* is one of the most tractable and comprehensively studied hymenopterans genetically^2^, owing to its overall ease of laboratory use, its short generation time (roughly ˜2 weeks), tolerance for inbreeding, and straightforward rearing. Similar to all other hymenopterans, *N. vitripennis* utilizes a haplodiploid sex determination system by which haploid males develop parthenogenetically from unfertilized eggs while diploid females develop from fertilized eggs^2^. Interestingly, this mode of sex determination makes *N. vitripennis* and other members of the clade vulnerable to manipulation by microbial and genetic parasites. For example, *Arsenophonus nasoniae*, a natural bacterial endosymbiont of *N. vitripennis*, effectively kills male progeny by manipulating key components of the mitotic machinery required specifically for early male embryonic development^3^. This male-killing results in significantly biased sex ratios favoring females, thereby benefiting the bacteria as they are transmitted solely from infected mother to offspring^4^. In addition to sex ratio-distorting bacteria, other genetic agents can influence the sex ratios of hymenopteran insects. For example, although the genome of *N. vitripennis* naturally harbors five chromosomes, some individuals have been discovered to also contain a sixth, supernumerary (B) chromosome termed paternal sex ratio (PSR)^5^. PSR is paternally transmitted through the sperm and acts by completely eliminating the haploid genome, thereby converting what should be diploid females into haploid PSR transmitting males, thereby making it a remarkable and potent selfish chromosome^5,6^. While progress has been made toward uncovering PSR-expressed transcripts^7^, the mechanism of action of this B chromosome in the *N. vitripennis* genome largely remains to be elucidated.

The last decade has experienced a rapid increase in the genetic toolkit to study the biology of *N. vitripennis* and its interesting interactions with bacterial symbionts and genetic parasites. For example, the availability of its high-resolution sequenced genome^8,9^, and several recent tissue-specific gene expression studies, together have provided a wealth of developmental gene expression information to be functionally analyzed^7,10,11^. Furthermore, methods to functionally disrupt gene expression relying on RNA interference (RNAi) by injecting *in vitro* transcribed dsRNA into either female pupae^12^ or larvae^13^ have advanced capabilities of performing reverse genetics on this organism. Altogether, these features have rendered *N. vitripennis* as a burgeoning model organism^13-16^ for studying complex genetic, cellular and developmental processes including venom production^17,18^, sex determination^19^, host symbiont interactions^3,20^, evolution and development of axis pattern formation^21-24^, and development of haplodiploidy^24^.

While *N. vitripennis* has many amenable experimental tools and resources described above, to date there have been no successful methods developed that allow for direct gene mutagenesis in this organism. This absence can, in part, be attributed to the difficulty in using previous gene disruption technologies, e.g. TALENs and ZNFs^25^, in addition to a lack of detailed published protocols for easily performing embryonic microinjection in *N. vitripennis.* To overcome these significant limitations, here we have employed the CRISPR-Cas9 (clustered regularly interspaced short palindromic repeats) gene editing system in *N. vitripennis.* As a part of this system we developed an effective method for pre-blastoderm stage embryonic microinjection in this organism. We report robust embryonic survival rates following embryo microinjection, and high mutagenesis rates of the conserved eye marker gene *cinnabar* in surviving CRISPR-Cas9 injected individuals. Overall, we demonstrate an efficient, effective, inexpensive, and straightforward CRISPR-Cas9 heritable gene disruption approach for *N. vitripennis*, and to our knowledge this study represents one of the first gene disruption-based techniques conducted in a hymenopteran insect.

## Results

### Development of an CRISPR/Cas9 embryo microinjection protocol

For delivery of CRISPR-based reagents we initially established efficient techniques for egg collection, pre-blastoderm stage embryo microinjection, and subsequent rearing and genetics, before proceeding. Briefly, as illustrated in figure 1, our techniques involved (i) permitting male and female adults to mate (˜4 days), (ii) supplying fresh host fly pupae *(Sarcophaga bullata)* to mated females for oviposition (˜45 minutes), (iii) carefully opening the parasitized host pupae to collect pre-blastoderm stage wasp embryos (˜15 minutes), (iv) aligning these embryos on sticky tape (˜15 minutes), (v) micro-injecting embryos with CRISPR/Cas9 components (˜15 minutes), (vi) carefully placing injected embryos back into the pre-stung hosts for proper development (˜15 minutes), (vii) and transferring the parasitized hosts harboring the CRISPR/Cas9 injected embryos into a humidified chamber with roughly 70% relative humidity to prevent dehydration of the embryos/host (˜15 minutes). These parasitized hosts were then incubated for roughly 14 days to permit the *N. vitripennis* embryos to complete development, and once the injected adults emerged from the host (viii), we isolated, mated and screened these individually for the presence of mutations (see Methods and Supplemental Methods for a comprehensive, step-by-step protocol). Remarkably, this entire protocol, from mating, to injecting, to hatching of injected individuals takes roughly 19 days for completion.

**Figure 1.**
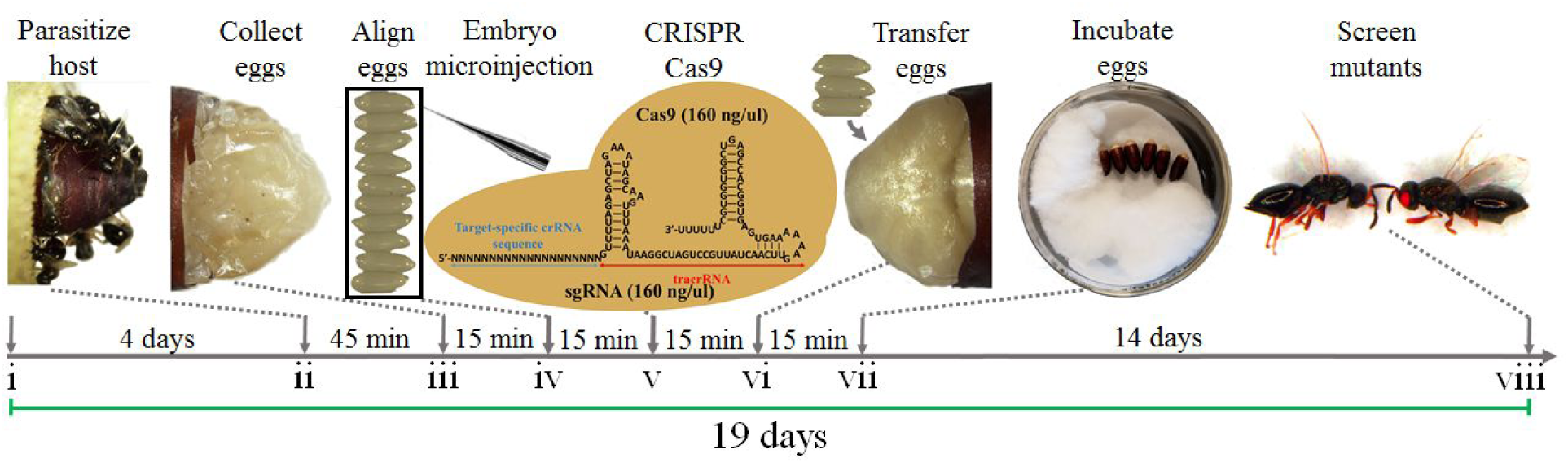
Schematic of *Nasonia vitripennis* embryo collection and CRISPR/Cas9 microinjections. Adult *Nasonia vitripennis* were mated for 4 days (i), then were supplied with a flesh fly host pupa, *Sarcophaga bullata*, for female parasitization for 45 minutes (ii). Embryos were then collected from the host (iii), aligned (iv), and injected with CRISPR/Cas9 components (v). Injected embryos were then gently placed back into the host (vi) for development (14 days) (vii), and when the adults emerged from the host they were subsequently screened for CRISPR/Cas9 induced mutations in target gene (viii). This entire procedure takes roughly 19 days to complete.

To initially test this injection protocol, we measured and compared the survival rates (to adulthood) of non-injected wasp embryos (*i.e.*, embryos removed from host, lined up on slide, then carefully placed back into host) to embryos injected with only purified water (*i.e.*, embryos removed from host, lined up on slide, injected with water, then carefully placed back into hosts). We found our survival rates to be quite robust for both non-injected embryos (92%), and for embryos injected with only water (76%).

### Identification of CRISPR/Cas9 target sites

To establish an efficient CRIPSR/Cas9 based genome editing platform for *N. vitripennis* we targeted the conserved dominant *cinnabar (cn)* gene (NV14284), which encodes for kynurenine hydroxylase, an enzyme involved in ommochrome biosynthesis^26^. Importantly, mutations in this gene result in distinct, scorable eye-color phenotypes when mutated in many organisms^27,28^, including *N. vitripennis* when silenced via larval RNAi^13^, thereby making it an optimal choice for the development and testing of a CRIPSR/Cas9 based gene mutagenesis technique in this organism. To disrupt this gene using CRISPR/Cas9, we designed several short guide RNAs (sgRNAs) to target either the third (sgRNA target sites 1 & 2) or the fourth (sgRNA target site 3) exons of the *cn* gene (Figure 2A). To define these specific exonic sgRNA genomic target sites we considered several factors. Firstly, we utilized available *N. vitripennis* transcriptional databases (http://www.vector.caltech.edu) to confirm *cn* RNA expression of the putative target regions^7,10^. Secondly, we searched both sense and antisense strands of the *cn* exon sequences of interest for the presence of the NGG protospacer-adjacent motifs (PAMs) utilizing CHOPCHOP v2 software^29^ and local sgRNA Cas9 package^30^. Thirdly, to minimize potential off-target effects, we confirmed specificity of our sgRNAs using publicly available bioinformatic tools^31^ and selected the most specific sgRNAs within our specified target region.

**Figure 2.**
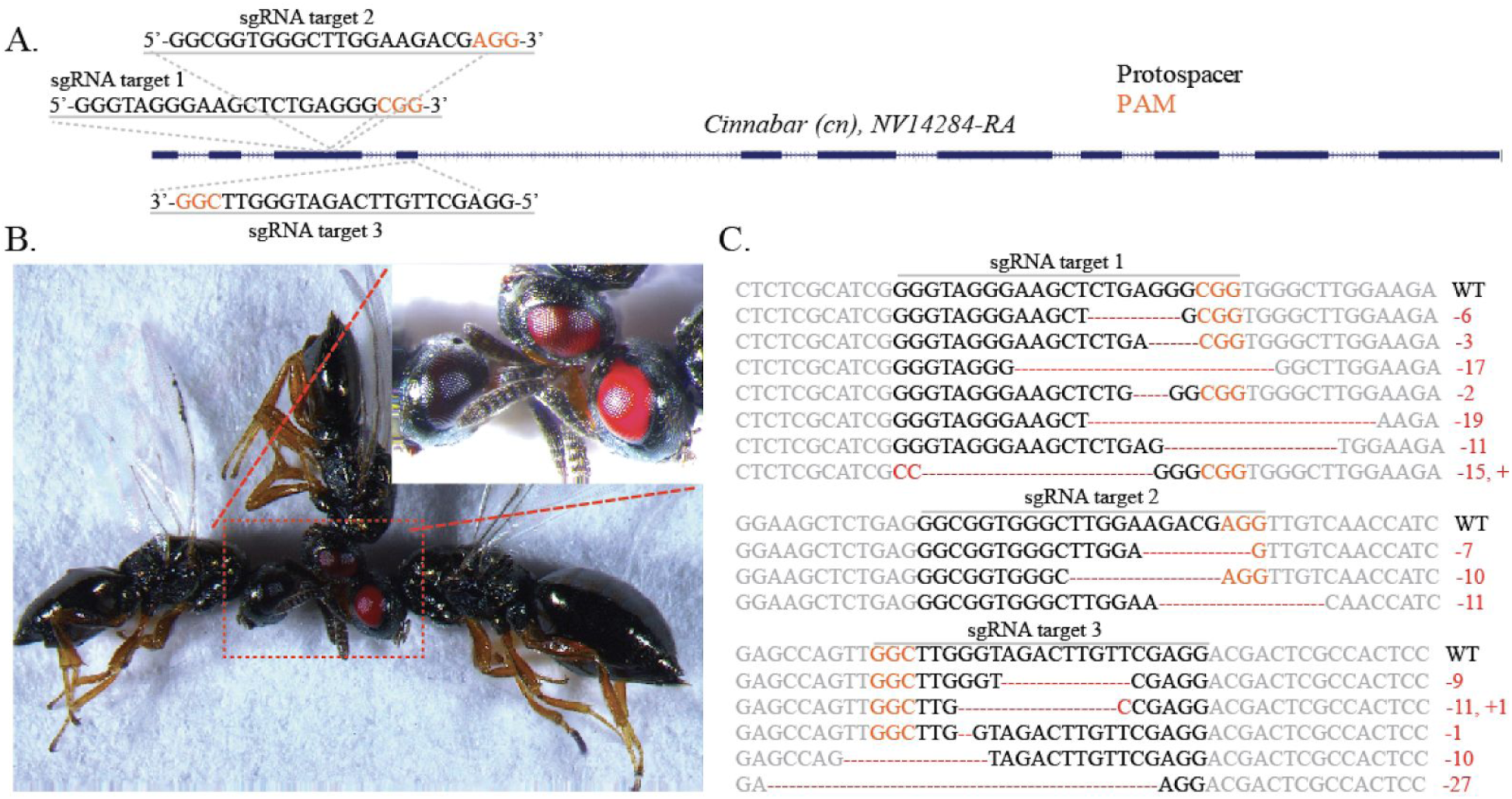
CRISPR/Cas9 target sites, mutant *cinnabar* phenotypes, and sequence disruption confirmations. Three independent sgRNAs were designed to target *cinnabar* in either exon 3 (sgRNA target 1 & 2) or exon 4 (sgRNA target 3) as depicted (A). Following embryo microinjection, surviving *cinnabar* mutant G0 adult wasps were readily observable with a light microscope by simply observing eye color phenotypes. Black eyes are wild-types, while red and bright red are mutants with different age (B). Many mutants for each sgRNA were established and deletions and insertions were readily detected via sequencing (C). PAM sequences (NGG) are indicated in orange, and *cn* gene disruptions resulting from insertions/deletions are indicated in red.

### Mutagenesis of the cinnabar gene is sgRNA/Cas9 dose dependent

To determine the optimal sgRNA/Cas9 concentrations for efficient disruption of *cn*, sgRNA-1 was chosen as a standard. We combined a variety of concentrations of sgRNA-1 (0, 20, 40, 80, 160, and 320 ng/ul) with the Cas9 protein (0, 20, 40, 80, 160, and 320 ng/ul) and found that the survival rate of the injected embryos, and the efficiency of mutagenesis mediated by CRISPR/Cas9, were dose-dependent (Table 1). These components also had an inverse relationship to each other; as the increased concentration of sgRNA and Cas9 protein lead to the increased proportion of red eye mutant adults (up to 60% of adult G0 survivors), the survival rate of injected eggs concomitantly decreased (Table 1). Therefore, we used the optimal combination of 160 ng/ul sgRNA and 160 ng/ul Cas9 protein as the working concentration for subsequent experiments.

**Table 1.** Effect of sgRNA and Cas9 protein concentration on *N. vitripennis* survival and mutagenesis

| sgRNA-1 | Cas9 | Total embryos | Adult Survivors |  |  | Mosaic (%) |  |  |
| --- | --- | --- | --- | --- | --- | --- | --- | --- |
|  |  |  | M | F | Total (%) | M (%) | F (%) | Both (%) |
| No injection | No injection | 100 | 66 | 26 | 92 (92) | 0 (0) | 0 (0) | 0 (0) |
| Water | Water | 100 | 44 | 32 | 76 (76) | 0 (0) | 0 (0) | 0 (0) |
| 20 (ng/ul) | 20 (ng/ul) | 100 | 34 | 34 | 68 (68) | 0 (0) | 0 (0) | 0 (0) |
| 40 | 40 | 100 | 30 | 32 | 62 (62) | 4 (13) | 0 (0) | 4 (6) |
| 80 | 80 | 100 | 24 | 22 | 46 (46) | 6 (25) | 0 (0) | 6 (13) |
| 160 | 160 | 100 | 16 | 22 | 38 (38) | 5 (31) | 3 (14) | 8 (21) |
| 320 | 320 | 100 | 10 | 10 | 20 (20) | 6 (60) | 6 (60) | 12 (60) |

To expand these studies and test other CRISPR/Cas9 target sites of *cn*, we injected an increased number of embryos (N=300) for each sgRNA/Cas9 combination, using our optimized sgRNA/Cas9 concentrations 160 ng/ul (Table 1), and had survival rates ranging from 22-27% of total embryos injected. From these injections, we discovered that 32% and 36% of injected survivor G0 *N. vitripennis* adults displayed the *cn* mutant phenotypes (*i.e.*, complete bilateral red eyes, Figure 2B) following microinjection with either sgRNA-1/Cas9, or sgRNA-3/Cas9 complexes, respectively (Table 2). However, lower mutagenesis efficiency (10%) was observed when sgRNA-2 was utilized, presumably resulting from inefficiency of sgRNA-3 (Table 2). Furthermore, in some instances we observed surviving G0 adults expressing a variegated (i.e., mottled) red/black eye phenotype (not shown), or in some cases, unilateral disruption (i.e., one complete black eye and one complete red eye in the same individual, not shown), as opposed to complete bilateral red eyes (mutant, Figure 2B) or complete bilateral black eyes (WT, Figure 2B), which we attributed to gene editing occurring in some nuclei at nuclear divisions past the first embryonic mitotic division (e.g. 2-nucleus embryo stage or later). Overall, these results strongly demonstrate the efficiency of the CRISPR/Cas9 system in *N. vitripennis* targeting multiple independent sites.

**Table 2.**
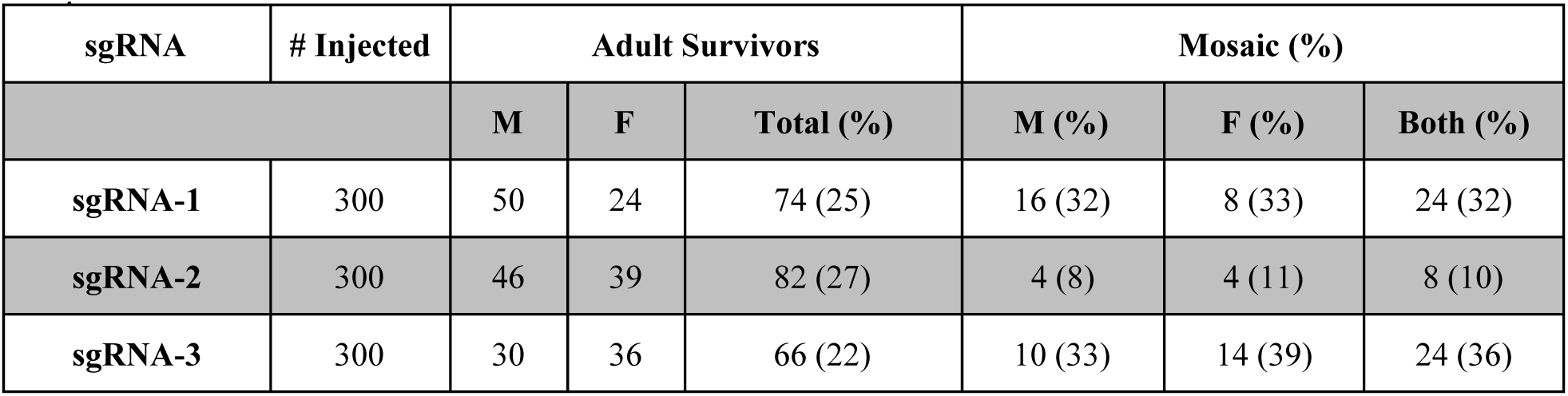
Summary of the injection and mutagenesis mediated by independent sgRNAs in *N. vitripennis*

| sgRNA | # Injected | Adult Survivors |  |  | Mosaic (%) |  |  |
| --- | --- | --- | --- | --- | --- | --- | --- |
|  |  | M | F | Total (%) | M (%) | F (%) | Both (%) |
| sgRNA-1 | 300 | 50 | 24 | 74 (25) | 16 (32) | 8 (33) | 24 (32) |
| sgRNA-2 | 300 | 46 | 39 | 82 (27) | 4 (8) | 4 (11) | 8 (10) |
| sgRNA-3 | 300 | 30 | 36 | 66 (22) | 10 (33) | 14 (39) | 24 (36) |

### Transmission of mutations to subsequent generations

Germline transmission of the CRISPR/Cas9 mutations to subsequent generations is essential for establishing stable mutant stocks. Given that all hymenopteran insects have haplodiploid sex determination with no heteromorphic sex chromosomes, and the widespread mode or reproduction is arrhenotoky, by which males develop from unfertilized eggs and are haploid, and females develop from fertilized eggs and are diploid, these factors had to be taken into consideration when designing the genetic crossing schemes to homozygose these mutant strains. Therefore, to test the germline transmission efficiency of the mutations generated by CRISPR/Cas9, and to establish homozygous mutant stocks, four crossing strategies were employed (Table 3). Overall, the results indicated that mutations are produced within the germline and transmitted to the subsequent generations with very high efficiency (e.g. 100% all male G1 offspring contained the mutant eye with crossing strategy D) and stable 100% mutant (male and female) producing lines could be produced by the G3 generation with the various crossing strategies. Together, these results indicated that the mutations had been efficiently transmitted into the germline and can be maintained in subsequent generations. Additionally, induced mutations can be obtained from either the G0 male or female parental direction.

**Table 3.** Summary of G1, G2 and G3 phenotypes of *N. vitripennis* with different crossing strategies

| Crossing strategy | G1 adult phenotype |  | G2 adult phenotype |  | G3 adult phenotype |  |
| --- | --- | --- | --- | --- | --- | --- |
|  | M<sup>w-</sup> (%) | F<sup>w-</sup> (%) | M<sup>w-</sup> (%) | F<sup>w-</sup> (%) | M<sup>w-</sup> (%) | F<sup>w-</sup> (%) |
| **A<sup>#</sup>** | 133 (92) | 31 (95) | 773 (100) | 336 (100) | 3741 (100) | 1159 (100) |
| **B<sup>†</sup>** | 0 (0) | 0 (0) | 71 (31) | 62 (17) | 659 (100) | 271 (100) |
| **C<sup>\*</sup>** | 12 (7) | 0 (0) | 93 (35) | 57 (22) | 1318 (100) | 577 (100) |
| **D<sup>%</sup>** | 52 (100) | 0 (0) | 0 (0) | 0 (0) | 1173 (24) | 227 (33) |

<sup>#</sup> G0 mutant ♂ X G0 mutant ♀, G1 mutant ♂ X G1 mutant ♀, G2 mutant ♂ X G2 mutant ♀
<sup>†</sup> G0 mutant ♂ X wild type ♀, G1 self-cross, G2 mutant ♂ X G2 mutant ♀
<sup>\*</sup> G0 mutant ♀ X wild type ♂, G1 self-cross, G2 mutant ♂ X G2 mutant ♀
<sup>%</sup> G0 mutant ♀ unmated, G1 mutant ♂ X wild type ♀, G2 mutant ♂ X G2 mutant ♀

Finally, to conclusively confirm the phenotypic defects described above were due to the genomic mutagenesis of the *cn*, genomic DNA was extracted from several independent mutant G2 lines and used as the template to PCR amplify the genomic DNA fragment containing the *cn* sgRNA target sites. The sequencing results confirmed the insertions/deletions in *cn* for all three sgRNA target sites tested (Figure 2C). Additionally, all sequenced lesions disrupted by the sgRNA target sequences generated deletions ranging from the loss of a single nucleotide to the loss of 27 nucleotides, and in some cases adding additional nucleotides around the targeted sites, in all cases disrupting gene function (Figure 2C).

## Discussion

Over the past decade or so, a number of important genetic, genomic, and cell biological studies have been conducted in the jewel wasp *N. vitripennis^19,32-35^*. These studies have been facilitated by the development of several important experimental resources including a high resolution genome sequence^8^, several genomewide transcriptional profiling studies^7,10^, procedures for performing embryonic *in situ* hybridizations to detect spatial patterns of mRNA expression^34^, and systemic, parental RNAi which can be used in certain tissue contexts to study gene function using reverse genetics^12,13^. Together these tools and others have progressively contributed to *N. vitripennis* becoming a preferred experimental system for hymenopteran-related biology. Notwithstanding these effective tools and resources, what has been lacking in *N. vitripennis* is a means for performing directed, heritable gene mutagenesis, which would facilitate efficient *in vivo* functional analysis of candidate genes in this species. To address this limitation, we tested whether the CRISPR/Cas9 system could be exploited as an effective gene editing platform in *N. vitripennis.* Overall our results demonstrate that the CRISPR/Cas9 system works efficiently in this organism; as a proof of principle we used this system to disrupt a conserved eye-pigmentation gene *cn*, utilizing several different sgRNAs with mutation rates up to 60%. Additionally, we found that these mutations were heritable, allowing us to homozygous them and establish stable mutant stocks.

Our study contains a few important caveats worth consideration. For example, we noticed that the efficiency of mutagenesis mediated by CRISPR/Cas9 and the survival rate of *N. vitripennis* injected embryos were sgRNA- and Cas9 protein-concentration dependent. Injected eggs with high concentrations of sgRNA and Cas9 combinations had higher mutagenesis rates but lower survival rates. A similar effect was also reported in other insects^36,37^, indicating that, on one hand, the concentration of injected sgRNA and Cas9 protein should be high enough to generate biallelic mutations to establish stable mutant populations; however, on the other hand, high concentration of sgRNA and Cas9 protein may cause toxic effects to the insects, thereby making it difficult to recover surviving mutant individuals. Our experiments suggest that an intermediate concentration of 160ng/ul for both the Cas9/sgRNA components achieves a moderate mutation rate while minimizing reduction of survivorship. We also noticed that the efficiency of cleavage is target site-dependent because each sgRNA we tested had a different cleavage rate (ranging between 10-36% of survivors). As others have reported, the chromatin environment around the target sites and sgRNAs sequence features have been identified as the major factors that affect the efficacy of CRISPR/Cas9 for any given target site^38^. As we have only targeted one gene, we have essentially assayed for only one chromatin environment that was conducive to gene editing. However, other genes may be affected negatively by different chromatin and sequence characteristics and, thus, variation in sgRNA targeting efficiency among targets differing in location across the genome is to be expected. Therefore, we recommend testing several sgRNAs for each gene to be targeted. Furthermore, in our study, mutations created in *cn* resulted in an easily scorable visible eye pigmentation phenotype which made screening of edited individuals straightforward. However, in reality many genes of interest, such as those involved in important cellular functions, will likely yield phenotypes such as sterility, lethality, or possibly even no visible phenotype when mutated, and will, therefore, require PCR-based genotyping. Additionally, in these cases screening and selection crosses will need to be revised in order to obtain the mutants and maintain them (*e.g.*, if disruption results in recessive lethal/sterile phenotypes the mutants must be maintained long term in a heterozygous state in the female sex and will require genotyping each generation).

Recently the CRISPR/Cas9 system has been demonstrated in the honey bee *Apis mellifera^39^*, and currently other groups are developing gene editing with this system in other hymenopterans. Here we have demonstrated that CRISPR/Cas9 should be widely applicable as a feasible means for gene editing in *N. vitripennis*, thereby further enhancing the tractability of this haplodiploid species as an insect system for the study of important biological questions that cannot be easily addressed in other hymenopterans that are less amenable to laboratory experimentation, or in other more traditional model organisms. While not tested here, this *N. vitripennis* CRISPR/Cas9 approach can be later expanded to test for integration of donor constructs via homology directed repair (HDR) following CRISPR mediated cleavage, similar to other species^40-43^. This modification will allow for site specific germline transformation and will further expand the *N. vitripennis* tool box, given that transgenesis still remains to be demonstrated, making it an even more useful model organism.

## Materials and methods

***Note - Information here provides a general overview of approaches and information on materials used, etc. A more detailed step-by-step protocol is supplied in the supplemental methods.***

### Production of sgRNAs

Linear double-stranded DNA templates for all sgRNAs were generated by template-free PCR with NEB Q5 high-fidelity DNA polymerase (catalog # M0491S) by combining primer pairs (sgRNA-1F & sgRNA-R) to make sgRNA-target-1, and combining primers paris (sgRNA-2F & sgRNA-R) to make sgRNA-target-2, and combining primers paris (sgRNA-3F & sgRNA-R) to make sgRNA-target-3. PCR reactions were heated to 98°C for 30 seconds, followed by 35 cycles of 98°C for 10 seconds, 58°C for 10 seconds, and 72°C for 10 seconds, then 72°C for 2 minutes. PCR products were purified with Beckman Coulter Ampure XP beads (catalog #A63880) according to the manufacturer protocol. Following PCR, sgRNAs were synthesized using the Ambion Megascript T7 *in vitro* transcription kit (catalog # AM1334, Life Technologies) according to the manufacturer’s protocols using 300ng of purified DNA template overnight at 37 °C. Following *in vitro* transcription, the sgRNAs were purified with MegaClear Kit (catalog #AM1908, Life Technologies) and diluted to 1000 ng/ul in nuclease-free water and stored in aliquots at −80°C. Recombinant Cas9 protein from *Streptococcus pyogenes* was obtained commercially (CP01, PNA Bio Inc) and diluted to 1000 ng/ul in nuclease-free water and stored in aliquots at −80°C. Immediately prior to injection, we combined the sgRNAs (at concentrations ranging from 20-320 ng/ul) with purified Cas9 protein (at concentrations ranging from 20-320 ng/ul) in purified water and pre-blastoderm embryonic microinjections were performed. All primer sequences can be found in table S1.

### Insect rearing, embryo collection, microinjection, transfer to hosts

*N. vitripennis* colonies were maintained in plastic cages (12 X 12 X 12 cm) and reared at 25 ± 1 °C with 30% humidity and a 12:12 (Light: Dark) photoperiod. Adults were fed with a 1:10 (v/v) honey/water solution that was provided in small droplets daily in a petri dish. Flesh fly pupa, *Sarcophaga bullata* (item number 144440) were ordered from www.carolina.com in batches of 100. To collect pre-blastoderm stage embryos, females and males were mated for at least 4 days. Following mating, we placed fresh *Sarcophaga bullata* pupae (hosts) into the cage to allow female wasps to parasitize the hosts for 45 minutes. Following parasitization, we carefully peeled off the puparium from the *Sarcophaga bullata* host pupae using forceps under a dissecting microscope and gently removed the recently laid exposed *N. vitripennis* embryos (<45 minutes old). We then quickly positioned these embryos onto a glass slide with double-sided sticky tape and injected the Cas9 protein and sgRNA mixtures into the germ cells located at the posterior of the *N. vitripennis* embryos. For microinjection consistency, we used a the Femtojet Express system (Eppendorf) with aluminosilicate glass filaments (Sutter Instrument). Following microinjection, we immediately placed the injected embryos back into pre-stung *Sarcophaga bullata* pupae with an ultra fine tip paintbrush, and incubated the embryos in a humidified chamber at 25°C until hatching.

### Cas9/gRNA-mediated mutation screens

Upon hatching, the mosaic phenotype in the G0 (injected wasps) was readily observed and assessed under microscope. Mutant individuals were isolated and mated using various crossing schemes to establish homozygous mutant stocks (table 3). To characterize the induced mutations, genomic DNA was extracted from individual wasps with the DNeasy blood & tissue kit (QIAGEN) following the manufacturer protocol. Target loci were amplified by PCR (using primers PCR-F and PCR-R), and the PCR product was analyzed via sequencing. Mutated alleles were identified by comparison with the wild-type sequence. All photographs were obtained using fluorescent stereo microscope (Leica M165FC). Primers used for PCR and sequencing are listed in table S1.

## Acknowledgements

This work was supported by generous University of California, Riverside (UCR) laboratory start-up funds to O.S.A, and a USDA National Institute of Food and Agriculture (NIFA) Hatch project (1009509) to O.S.A, and an NSF CAREER award (NSF1451839) to P.M.F.

## Disclosure

The authors declare no competing financial interests.

## Supplemental Protocol

### Goals

Heritable manipulation of the *N. vitripennis* genome through precise double-strand breaks at targeted gene sites via CRISPR-Cas9. This protocol was based on several CRISPR/Cas9 protocols developed in other species including *Drosophila melanogaster^1^*, *Aedes aegypti^2^*, and *Danio rerio^3^.*

- Efficient generation of CRISPR/Cas9 reagents: sgRNAs (Cas9 protein is commercially purchased).
- Microinjection of CRISPR/Cas9 components into N. vitripennis pre-blastoderm embryos.
- Isolation of mutant alleles generated by CRISPR/Cas9 mediated non-homologous end joining (NHEJ).
- Generation of stable mutant *N. vitripennis* strains through genetic crossing.

### General Considerations

- It is very important to work under RNAse free conditions when producing or working with sgRNAs. Be sure to use nuclease-free consumables including filter tips and microfuge tubes. Also, thoroughly clean work area, microinjection apparatus, gloves, and pipettes with RNaseZap (Ambion) before conducting experiments.
- Once sgRNAs are produced, immediately mix these reagents with Cas9 protein at the final desired injection concentrations and make small 5-10ul aliquots. Store these ready-to-inject final mixtures at −80°C until needed. The goal here is to avoid excess freeze-thaw-cycles for both the sgRNAs and the Cas9 protein as much as possible.
- Given that not all sgRNAs function efficiently, and specificity and activity are unpredictable, we recommend designing multiple sgRNAs for each target gene to increase probability of generating desired modifications of target genes.
- To collect enough *N. vitripennis* eggs for microinjection, it is important to expand stock wasp colonies to sufficient numbers. We recommend to establish 2-3 colonies with 200-500 wasps per colony.

### Selected Kits and Reagents

- *Streptococcus pyogenes* Cas9 recombinant protein, PNA Bio (catalog # CP01)
- Flesh fly pupa, *Sarcophaga bullata*, www.carolina.com (catalog# 144440)
- Beckman Coulter Agencourt Ampure XP beads (catalog #A63880)
- Zymo Research DNA Clean and Concentrator kit (catalog #D4005)
- NEB Q5 High-Fidelity DNA polymerase (catalog # M0491S) - PCR of sgRNA
- Ambion MegaScript T7 (catalog #AM1334) – sgRNA *in vitro* transcription kit
- Ambion MegaClear Kit (catalog #AM1908) – Purification of *in vitro* transcribed sgRNA
- Agilent Bioanalyzer 2100 and RNA 6000 Nano Kit (catalog #5067-1511)
- Ambion RNaseZap (catalog # AM9780) – removal of RNAse’s from work area.
- DNeasy blood & tissue kit (QIAGEN, catalog # 69506)- genomic DNA isolation
- dNTP solution mix, 25 mM each (Enzymatics, catalog # N205)
- UltraPure DNase/RNase-free distilled water (Life Technologies, catalog # 10977-023)
- Double-sided sticky tape
- 10% (vol/vol) sucrose solution
- Microloader Tips for Filling Femtotips (Eppendorf, catalog # 930001007)
- sgRNA-F and sgRNA-R primers from IDT as PAGE-purified oligos

### Equipment

- Sutter instruments Microelectrode Beveler (Model # BV10)
- World Precision Instruments (WPI) micromanipulator (Model # Kite R)
- Eppendorf Femtojet Express programmable microinjector
- Sutter micropipette puller (P-1000, and P-2000)
- Olympus SZ51 -screening microscope
- Leica DM-750 -microinjection microscope
- Several bugdorm-41515 (L17.5 x W17.5 x H17.5 cm) cages
- Microinjection needles
  - Borosilicate glass capillary tubing 1 mm (outside diameter) × 0.58 mm (inner diameter) (Sutter Instrument, catalog # BF100-58-10)
  - Quartz glass capillary tubing 1 mm (outside diameter) × 0.70 mm (inner diameter) (Sutter Instrument, catalog # QF100-70-10)
  - Aluminosilicate glass capillary tubing 1 mm (outside diameter) × 0.64 mm (inner diameter) (Sutter Instrument, catalog # AF100-64-10)
- PCR Machine with heated lid (BIO-RAD, T100^TM^)

### Genomic DNA target site design and selection criteria

The revolutionary CRISPR/Cas9 gene editing system relies on the target sequence encoded by an engineered sgRNA to guide the Cas9 nuclease to a desired genomic target location, allowing for base-pairing interactions between the sgRNA/Cas9 complex and the complementary genomic DNA sequence, thereby resulting in subsequent Cas9 mediated cleavage of the specified genomic target sequence. The characterized recognition sequence for the *Streptococcus pyogenes* Cas9 protein relies on the presence of a protospacer associated motif, or PAM, to be located immediately adjacent to the desired genomic target sequence^4-6^. The PAM sequence is NGG, which is located in the genome directly downstream of the desired 20bp genomic target sequence, taking the form N20-NGG. Importantly, the PAM sequence is not included in the either the sgRNA template DNA, or the *in vitro* transcribed sgRNA. To define putative sgRNA genomic target sites we suggest to consider several factors. First, confirming transcriptional expression of the target region, and looking for conservation between other species, will help define putative *N. vitripennis* genomic target regions that have a higher probability of being necessary for gene function (assuming the goal is to disrupt gene function). To do this we recommend using both available *N. vitripennis* transcriptional databases and simple NCBI-BLAST searches www.vector.caltech.edu.^7,8^. Secondly, once general target regions are defined, the putative sgRNA target sites can be identified by simply scanning both the sense and antisense strands for the presence of the NGG-PAMs either manually by eye, or by utilizing available software such as CHOPCHOP v2^9^, and/or local sgRNA Cas9 packages^10^. Finally, to minimize potential off-target effects, we recommend confirming specificity of the sgRNAs using publicly available bioinformatic tools^11^ and selecting the most specific sgRNAs within the specified target regions with the least potential off-target binding sites. It should be noted that even if the chosen sgRNA target sequences fulfill all of the above requirements, sgRNA specificity and activity is unpredictable. Therefore, we recommend that multiple different sgRNAs are designed and co-injected in order to target a given gene of interest.

Shown below is an example of a genomic target region depicted in figure 1 with three chosen sgRNA target sequences. The primary genomic target sequences are highlighted in green, while the PAMs are highlighted in blue (NGG).

- Target # 1: ACATTACATCGGAATCGTAC CGG
- Target # 2: GGATCCTCATGACATTACAT CGG
- Target # 3: GATTCCGATGTAATGTCATG AGG

Importantly, the first two nucleotides transcribed by the T7 RNA polymerase should ideally be “GG,” and therefore, it is important that the *in vitro* transcribed sgRNA sequences beings with these two nucleotides. Therefore, if perhaps the chosen sgRNA sequences (green) do not begin with “GG,” these nucleotides can be added onto the 5’ end of the target sequence. For example, for the target sequences highlighted in figure 1, the following bases (pink) would be added to ensure robust *in vitro* transcription by the T7 RNA polymerase. For target # 1, two GG’s would need to be added, for target # 2 no extra GG’s would need to be added since it already begins with a GG, and for target # 3 one G would need to be added to ensure the first two nucleotides (<u>underlined</u>) begin with “GG.”

- Target # 1: <u>GG</u> ACATTACATCGGAATCGTACCGG
- Target # 2: <u>GG</u>ATCCTCATGACATTACATCGG
- Target # 3: <u>GG</u>ATTCCGATGTAATGTCATGAGG

### sgRNA DNA template generation

The first step in sgRNA production is to produce the template DNA to be used for *in vitro* transcription. This DNA is generated by template-free PCR using two primers that anneal to each other via complementary sequences (**<u>bold</u> <u>and</u> <u>underlined</u>**). These primers can be ordered from IDT as PAGE-purified oligos.

<u>sgRNA-R</u> (Table S1) - This is a universal reverse primer that can be used to generate all sgRNA targets containing the sgRNA backbone sequence.

- 5’-AAAAGCACCGACTCGGTGCCACTTTTTCAAGTTGATAACGGACTAGCCTTA TTTTAACTT**<u>GCTATTTCTAGCTCTAAAAC</u>**-3’

<u>sgRNA-F</u> - This primer contains the T7 promoter upstream (orange) of the target sequence.

- 5’–GAAATTAATACGACTCACTATA GG**N**^20^ **<u>GTTTTAGAGCTAGAAATAGC</u>**–3’

Note: The GG**N^20^** is a generic sequence that includes the 20 nucleotide user-defined genomic target sequence (**N^20^**), and the “GG” sequence necessary for in *vitro* T7 RNA polymerase transcription, but does **NOT** include the PAM sequence.

1. The first step in sgRNA production is to setup a template-free PCR using sgRNA-R and sgRNA-F primers to produce the linear dsDNA templates that will be used for *in vitro* transcription reactions. We prefer to use NEB’s Q5 high-fidelity DNA polymerase, however other DNA polymerases should also work. Below is the PCR reaction we recommend.

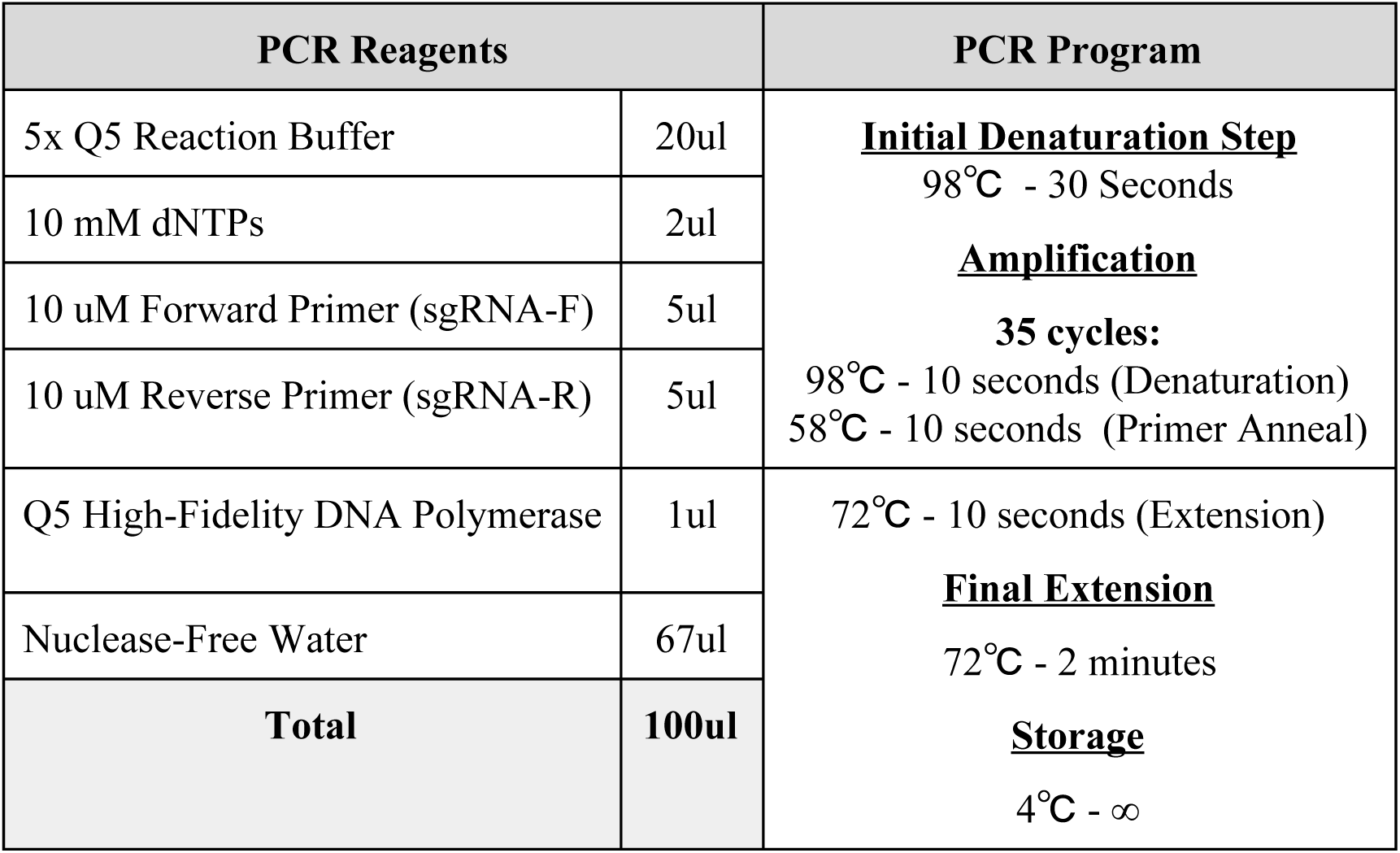
2. Following the PCR reaction, run 3ul of the PCR product on a DNA agarose gel (2%) to confirm the PCR reaction amplified the desired DNA and the size (band ˜122bp in size) is as expected.
3. The PCR products are then purified with Beckman Coulter Ampure XP beads (catalog #A63880) according to the manufacturer protocol. Note - instead of the Ampure XP beads, we have also used the Zymo DNA clean and Concentrator kit at this step and that also effective.

| PCR Reagents |  | PCR Program |
| --- | --- | --- |
| 5x Q5 Reaction Buffer | 20ul | **<u>Initial Denaturation Step</u>**<br>98°C - 30 Seconds<br>**<u>Amplification</u>**<br>**35 cycles:**<br>98°C - 10 seconds (Denaturation)<br>58°C - 10 seconds (Primer Anneal) |
| 10 mM dNTPs | 2ul |  |
| 10 uM Forward Primer (sgRNA-F) | 5ul |  |
| 10 uM Reverse Primer (sgRNA-R) | 5ul |  |
| Q5 High-Fidelity DNA Polymerase | 1ul | 72°C - 10 seconds (Extension)<br><u>**Final Extension**</u><br>72°C - 2 minutes<br><u>**Storage**</u><br>4°C - ∞ |
| Nuclease-Free Water | 67ul |  |
| **Total** | **100ul** |  |

Simplified Beckman Coulter Ampure XP beads protocol
a. Gently shake the Beckman Coulter Ampure XP bottle to resuspend the magnetic particles.
b. Add 1.8x volume of Ampure XP beads to the PCR reaction in a 1.5 ml microcentrifuge tube.
c. Mix reagents and PCR reaction thoroughly by pipetting up and down.
d. Incubate for 5 min at room temperature for maximum recovery.
e. Place the tube on a magnetic rack and wait until the liquid is clear to capture the beads. Carefully remove supernatant.
f. Keep the tube on the magnetic rack and add 300 ul freshly made 80% ethanol (make with nuclease-free water) to wash the beads.
g. Incubate the tube at room temperature and wait for 1 min.
h. Carefully remove supernatant.
i. Repeat steps f-h again.
j. Air dry the beads at room temperature for 5 min.
k. Remove the tube form the magnetic rack.
l. Resuspend the beads in 20 ul of nuclease-free water.
m. Incubate the tube at room temperature for 10 min to elute DNA from the beads.
n. Place the tube back to the magnetic rack until the liquid is clear.
o. Transfer supernatant to a new tube.

Simplified Zymo DNA clean and Concentrator kit protocol
a. In a 1.5 ml tube, add 5 volumes of DNA Binding Buffer to each volume of PCR reaction.
b. Mix reagent and PCR reaction thoroughly by pipetting.
c. Transfer mixture to a Zymo-Spin™ Column in a collection tube.
d. Centrifuge for 30 seconds (11000 x g) and discard the flow-through.
e. Add 200 ul DNA wash buffer to the column, and centrifuge for 30 seconds (11000 x g), discard the flow-through.
f. Repeat the step e again.
g. Add 20 ul nuclease-free water directly to the column matrix.
h. Incubate at room temperature for 1 min.
i. Transfer the column to a new 1.5 tube and centrifuge for 1 min (11000 x g) to elute the DNA.
4. Following purification, measure the purity and concentration of the purified DNA template using a nanodrop. We aim to have a concentration of over 100ng/ul to ensure enough template for the *in vitro* transcription reaction.

## sgRNA production by *in vitro* transcription

To produce the sgRNAs, we use the Ambion MegaScript T7 *in vitro* transcription kit and followed the manufacturer's protocol.

1. Briefly, we thaw and mix thoroughly the ribonucleotides (keep on ice) and reaction buffer (keep at room temperature), then add all reagents to a PCR tube in the following order.

2. Following *in vitro* transcription, 1ul of Turbo DNAse should be added to the reaction and incubated @ 37°**C** for 15 minutes to remove the template DNA from the reaction. The sgRNAs can then be purified with Ambion MegaClear Kit following the manufacturer protocol.

| ***In vitro* Transcription Reagents** | **Volume** | **Reaction** |
| --- | --- | --- |
| Nuclease free water | To 20uL | 2 hours (minimum) to overnight (12-16 hours maximum) @ 37°C. Note - we recommend overnight incubation here.<br>Add 1ul of Turbo DNase (mix well) and incubate @ 37°C for 15 minutes. |
| Free ribonucleotides (ATP, CTP, GTP, UTP) | 2ul each (8ul total) |  |
| 10x Reaction Buffer | 2ul |  |
| sgRNA PCR template | 300ng total (~3ul) |  |
| T7 Enzyme | 2ul |  |
| **Total** | **20ul** |  |

Simplified Ambion MegaClear protocol
a. In a 1.5 ml tube, bring the RNA sample to 100 ul with the elution solution. Mix gently but thoroughly by pipetting.
b. Add 350 ul of binding solution concentrate to the sample. Mix gently but thoroughly by pipetting.
c. Add 250 ul of 100% ethanol to the sample. Mix gently but thoroughly by pipetting.
d. Pipet the RNA mixture above onto the filter cartridge and centrifuge for 1 min at RCF 13000 x g.
e. Discard the flow-through.
f. Wash with 500 ul wash solution, discard the flow-through.
g. Repeat the step f.
h. After discarding the wash solution, centrifuge the filter cartridge for 1 min at RCF 13000 x g.
i. Place the filter cartridge into a new 1.5 ml tube.
j. Add 50 ul of nuclease-free water to the center of the filter cartridge.
k. Close the cap of the tube and incubate at 70°C for 10 min.
l. Centrifuge (13000 x g) for 1 min at room temperature to elute RNA.
3. The final concentration should be measured using a nanodrop, and quality can be measured with an Agilent Bioanalyzer confirming that sgRNA appears as a single band without any degradation products.
4. sgRNAs can then be diluted to 1000 ng/ul in nuclease-free water and stored in aliquots @ −80°C. We generally produce roughly 5-100ug of sgRNA from this reaction depending on the template DNA quality.

## Preparation of sgRNA/Cas9 mixtures for microinjection

Before microinjection the purified recombinant Cas9 protein from *Streptococcus pyogenes* should be obtained commercially (CP01, PNA Bio Inc) and diluted to 1000ng/ul using UltraPure DNase/RNase-free distilled nuclease free water and stored @ −80°C.

- This stock Cas9 protein solution should be diluted with nuclease free water and mixed with the purified sgRNAs at various concentrations (20-320ng/ul) in small 5-10ul aliquots.
- These ready-to-inject final mixtures can be stored at −80C until needed. The goal here is to avoid excess freeze-thaw-cycles for both the sgRNAs and the Cas9 protein as much as possible.
- For *N. vitripennis*, we found the optimal concentrations for both the Cas9 protein and purified sgRNAs to be 160ng/ul for each component.
- To prepare these mixtures thaw and mix both components in UltraPure DNase/RNase-free distilled nuclease free water on ice, and maintain these mixtures on ice while performing injections.

## Preparation of needles for *N. vitripennis* embryo microinjection

For effective penetration and microinjection into *N. vitripennis* eggs, we experimented with several types of capillary glass needles with filament including Quartz, Aluminosilicate and Borosilicate types. The quality of needles is critical for avoiding breakage/clogging during injection, embryo survival and transformation efficiency. For each of these glass types we developed effective protocols to pull these needles on different Sutter micropipette pullers (P-1000, and P-2000) to enable the needles to have a desired hypodermic-like long tip that we found effective for *N. vitripennis* embryo microinjection. The parameters (filament, velocity, delay, pull, pressure) for the different types of capillary glass needles are listed in the following table. While all three types of needles were effective for *N. vitripennis* injections, we prefered the Aluminosilicate capillary glass needles, because the Quartz capillary glass needles were too expensive, and the Borosilicate capillary glass needles were a bit too soft and clogged easily.

| Capillary glass needle type | Sutter Needle Puller model | Heat | Filament | Velocity | Delay | Pull | Pressure |
| --- | --- | --- | --- | --- | --- | --- | --- |
| Quartz | P-2000 | 750 | 4 | 40 | 150 | 165 | - |
| Aluminosilicate | P-1000 | 605 | - | 130 | 80 | 70 | 500 |
| Borosilicate | P-1000 | 450 | - | 130 | 80 | 70 | 500 |

## *N. vitripennis* pre-blastoderm stage embryos collection and alignment

1. Before collecting embryos, it is important to expand *N. vitripennis* colonies and set up several (3-4) bugdorm-41515 (L17.5 x W17.5 x H17.5 cm) cages with roughly 200-500 adult wasps in each cage (figure 2). This will ensure enough eggs are laid on demand for microinjection.
2. Make sure the wasps are healthy, and well fed, by freshly providing small droplets of 1:10 (v/v) honey/water solution daily, and removing old honey/water solution. Maintain *N. vitripennis* colonies at 25 ± 1 °C with 30% relative humidity and a 12:12 (Light: Dark) photoperiod.
3. Allow the females and males to freely mate for at least 4 days, prior to injection, and keep them completely starved of hosts to ensure females lay eggs when needed.
4. When ready to collect embryos, place a few (2-3) fresh *Sarcophaga bullata* pupae into the cage with the gravid wasps. Importantly, use a foam stopper to only expose only about 0.5 cm of the hosts for parasitization to ensure that the embryos are laid in a concentrated manner at the posterior end of the host for rapid egg collection (figure 3). Alternatively, a 1-ml pipette tip cut ˜0.5 cm from the end can also be used to restrict egg laying on host as described previously^12^.
5. Allow female wasps to parasitize (oviposit embryos) the host for roughly 30 minutes at 25^0^**C**. Then remove the host and replace with a new host, every 15 minutes, to ensure sufficient eggs for continuous injection. Note - it is very important that the embryos are as young as possible, ideally within the first hour of being oviposited, to ensure that they are in the pre-blastoderm stage. Old embryos (>1.0 hour) should not be injected.
6. To collect embryos, remove parasitized hosts from the foam stopper. Under a dissecting microscope, carefully peel off the posterior end of the puparium that was exposed to the wasps using forceps. Embryos will be resting on the surface of the host pupa (figure 4). Carefully remove embryos from host, using a fine-tip wet paintbrush, ensuring not to burst the soft pupal skin inside the host.
7. Transfer embryos one-by-one to double-sided sticky tape (fixed to a glass slide). Using a wet paintbrush orient the eggs one-by-one in a row so the posterior end (more narrow end) is pointing in the same direction for each egg (figure 5). Note - we found embryo survival rates to be greater if we did not cover eggs with halocarbon oil during injection as is done for *Drosophila melanogaster* microinjection^13^. Since oil is not used, it is important to keep the embryos moist during the injection period by regularly adding water using the paintbrush. The amount of water on the brush is key to move embryos around and align with ease. Too much water results in embryos floating and too little water makes them difficult to move around. To adjust the degree of moisture, dip the tip of the brush in water then lightly touch the tip of the brush to a dry kimwipe.
8. Ideally this protocol is most effective if one person is continuously collecting and lining up eggs, while the other person is injecting the CRISPR/Cas9 components.

## CRISPR/Cas9 embryo Microinjection

1. Break the closed tip of the needle by either rubbing the needle to the edge of the slide, or by using a microelectrode beveler (Sutter Instrument).
2. Load the needle with 2ul of injection mixture using Microloader Tips for Filling Femtotips.
3. One-by-one inject, 1-5pl of injection mixture (about 2 −10% of the egg’s volume) into the cytoplasm from the posterior end of each egg. We use a femtojet express to control for the injection volume.
4. Inject ˜40 eggs at a time (should take roughly 10 minutes) then stop and transfer injected eggs into a host then continue injecting again using a fresh newly laid batch of eggs.

## Transferring embryos back to the hosts

1. Following microinjection, transfer injected embryos back into a pre-stung *Sarcophaga bullata* pupae using a fine-tip paintbrush (figure 6). *N. vitripennis* larva utilize the host pupa as a food source to complete larval development and to our knowledge there is currently no available artificial diet that can be used.
2. Very important - be sure to **only** transfer eggs back into a **pre-stung host**, otherwise embryos will not survive. When a female wasp stings a host, she uses her ovipositor to bore a hole in the host puparium to inject venom which causes arrest of the pupal development, allowing the *N. vitripennis* larvae to consume the host. Without the venom, the host will survive and the *N. vitripennis* larvae will not be able to consume the host.
3. To ensure a host has already been stung, find a host with embryos in it, then scrape all the embryos off and use it as the host. Also, to avoid overcrowding, only place about 40 injected embryos or less per host.
4. Incubate hosts harboring transferred injected eggs in a moist humidified chamber (e.g. petri dish with cotton balls moist with water) at 25°C until hatching (roughly 1-2 days). Importantly, hosts can be left with a peeled off puparium and the *N. vitripennis* eggs will develop normally so long as they are incubated in a humidified chamber (petri dish with damp filter paper and cotton balls) with roughly 70% relative humidity (figure 7).
5. Monitor the embryos, the hatched *N vitripennis* larvae, and the host daily. Remove any dead *N. vitripennis* larvae, and if the host becomes infected with bacteria or dies (turns to the gray or dark color and has a foul smell) transfer the larvae to a fresh pre-stung host.

## Screening for modification and Genetics

1. After roughly 8 days the injected embryos will begin to pupate. Once they pupate they will no longer consume food (i.e. blowfly host) and can be removed from the host.
2. Remove each *N. vitripennis* pupae from the host, and place in an individual 1.5ml eppendorf tube until hatching. This will ensure that the hatched females will be virgin and will not mate until desired.
3. Upon hatching, if disrupting a visual marker gene (e.g. *cinnabar)* then the mutant phenotype should be readily visible. If disrupting a non-visible marker gene, then every surviving G0 (injected individual) should be mated with wildtype individually, and given a separate host to produce its own colony. Importantly, similar to *Drosophila melanogaster*, *N. vitripennis* can be immobilized by exposure to CO_2_ allowing for straightforward manipulation.
4. Once the injected G0 males and females and have successfully mated, and produced progeny, the G0’s can be sacrificed and genomic DNA should be extracted using the DNeasy blood & tissue kit for every individual.

Simplified DNeasy blood & tissue kit Protocol
a. Place the sample into a sterile 1.5 ml microcentrifuge tube.
b. Add 180 ul buffer ATL and 20 ul proteinase K, mix by vortexing 10-15 seconds.
c. Incubate the sample overnight at 56 °C until completely lysed.
d. Add 200ul buffer AL. Mix thoroughly by vortexing.
e. Add 200ul ethanol (96%-100%). Mix thoroughly by vortexing.
f. Pipet the mixture into a DNeasy mini spin column placed in a 2 ml collection tube.
g. Centrifuge at 8000 x g for 1 min.
h. Discard the flow-through and collection tube. Place the spin column in a new 2 ml collection tube.
i. Add 500 ul buffer AW1. Centrifuge for 1 min at 8000 x g.
j. Discard the flow-through and collection tube. Place the spin column in a new 2 ml collection tube.
k. Add 500 ul buffer AW2, and centrifuge for 3 min at 20000 x g.
l. Discard the flow-through and collection tube. Transfer the spin column to a new 1.5 ml tube.
m. Add 30 ul buffer AE to the center of the spin column membrane.
n. Incubate for 1 min at room temperature.
o. Centrifuge for 1 min at 8000 x g.
5. The presence of mutations can be determined by PCR amplifying/sequencing the genomic target region.
6. Colonies that have mutations as determined by sequencing should be continued, while colonies that were established with non-mutant G0’s should be discarded.
7. Importantly, unmated females will give rise to 100% haploid male broods, so therefore a mutant unmated female can give rise to large number of knockout males that can be used for subsequent analysis.

**Table S1.**
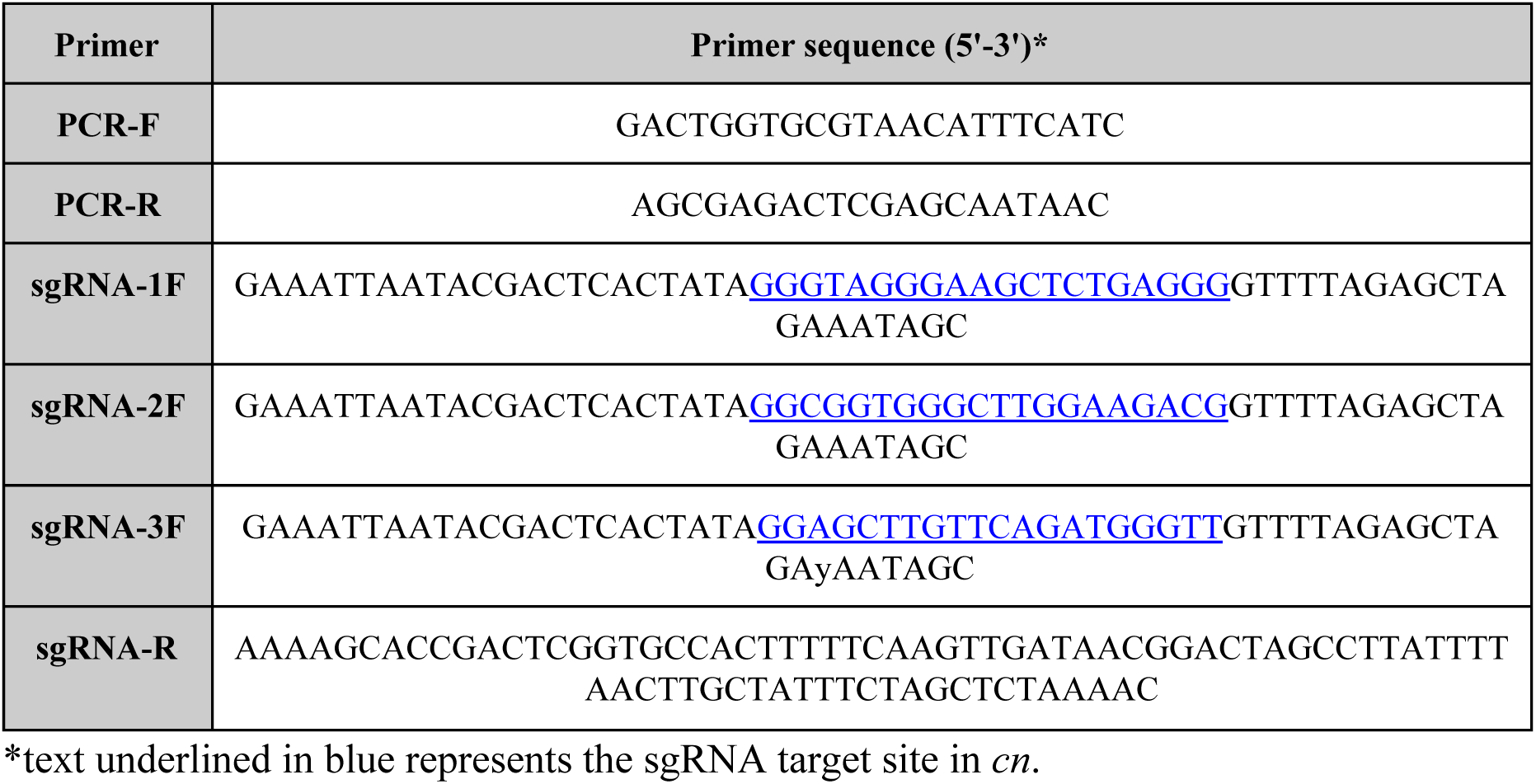
Primers used in this study.

**Figure 1.**
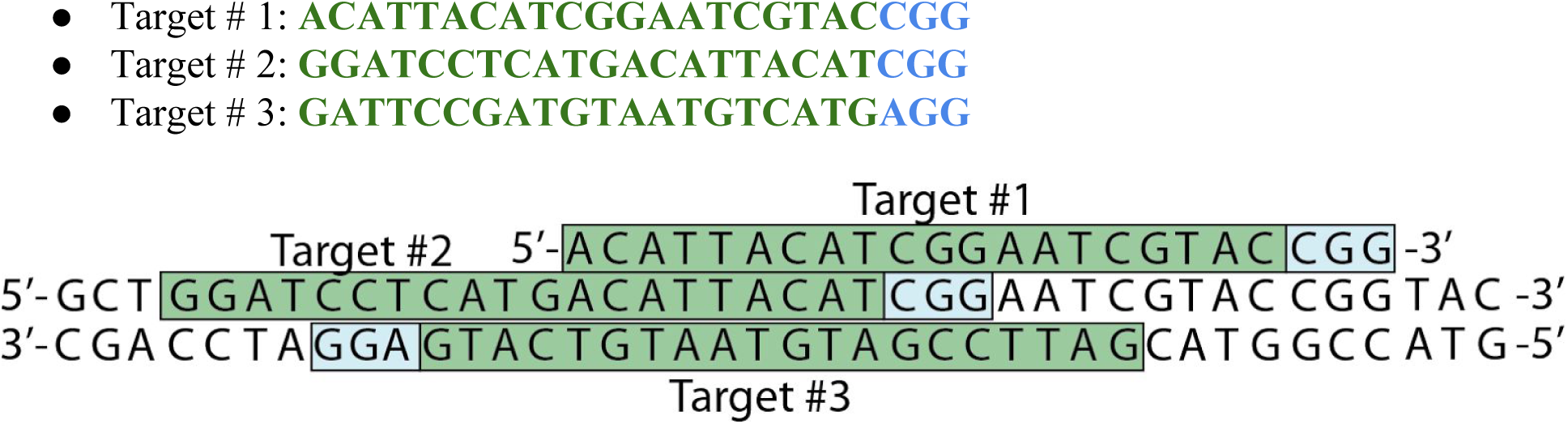
A genomic DNA example target region with target sites (#1-3) highlighted in green and the PAM sequences are highlighted in blue. Targets sites #1&2 are located on the top strand, while target site # 3 is located on the bottom strand.

**Figure 2.**
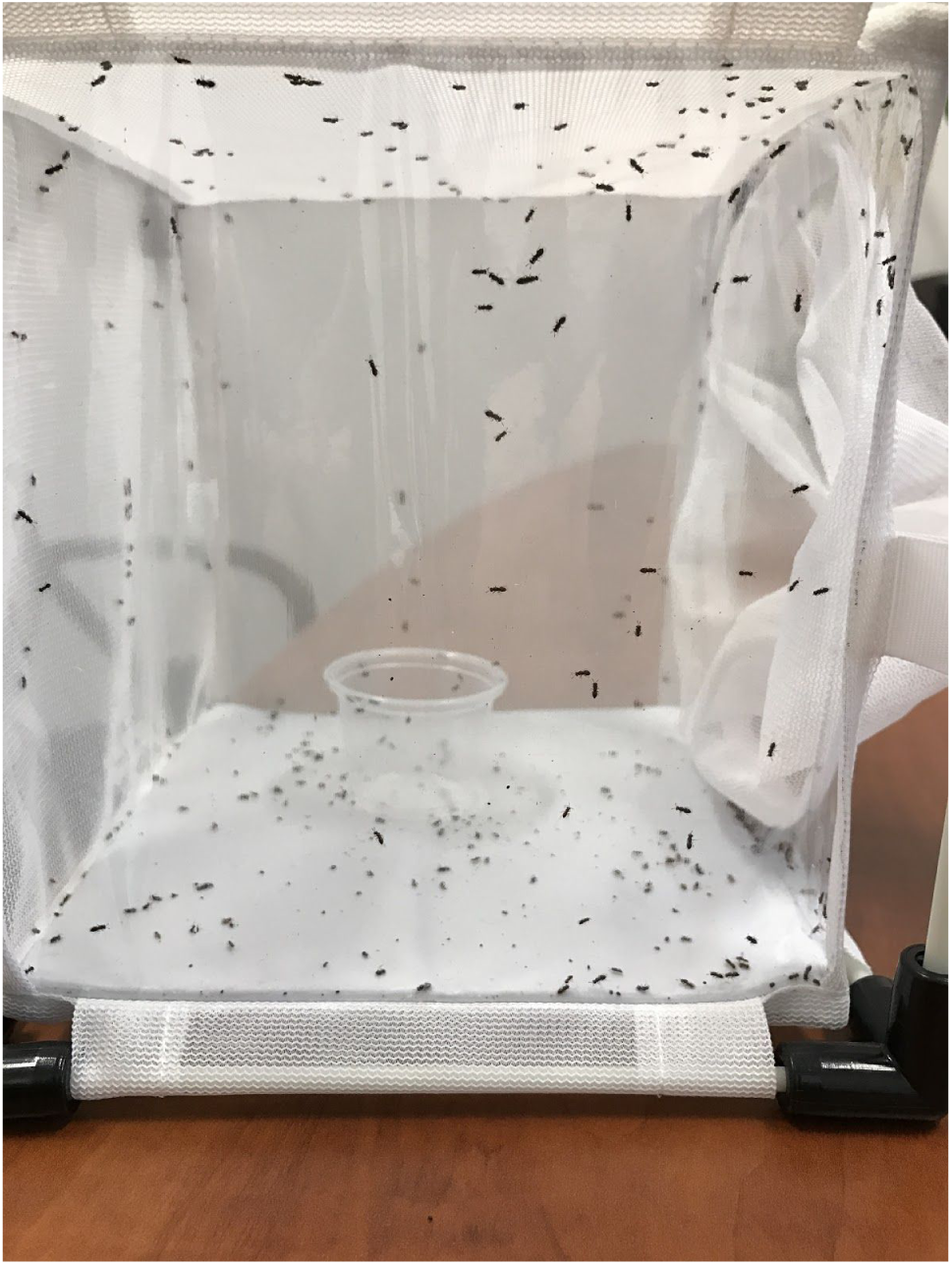
*N. vitripennis* colony in a bugdorm-41515 (L17.5 x W17.5 x H17.5 cm) with roughly 200-300 adult wasps.

**Figure 3.**
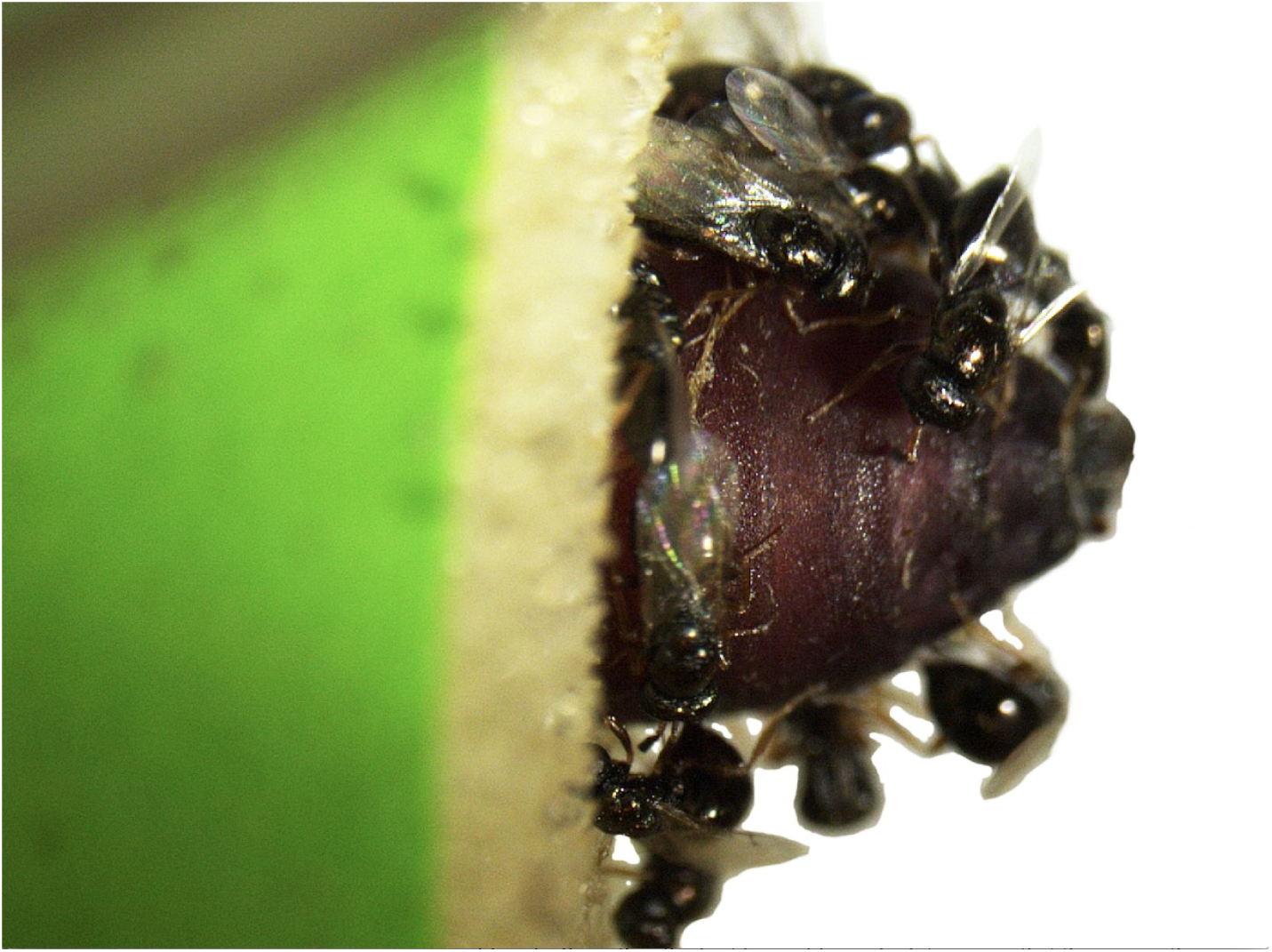
Gravid *N. vitripennis* females parasitizing 0.5cm of the posterior end of a *Sarcophaga bullata* pupae. Importantly, the pupae, is placed inside of a foam stopper (green) to ensure embryos are laid in a concentrated manner at the posterior end.

**Figure 4.**
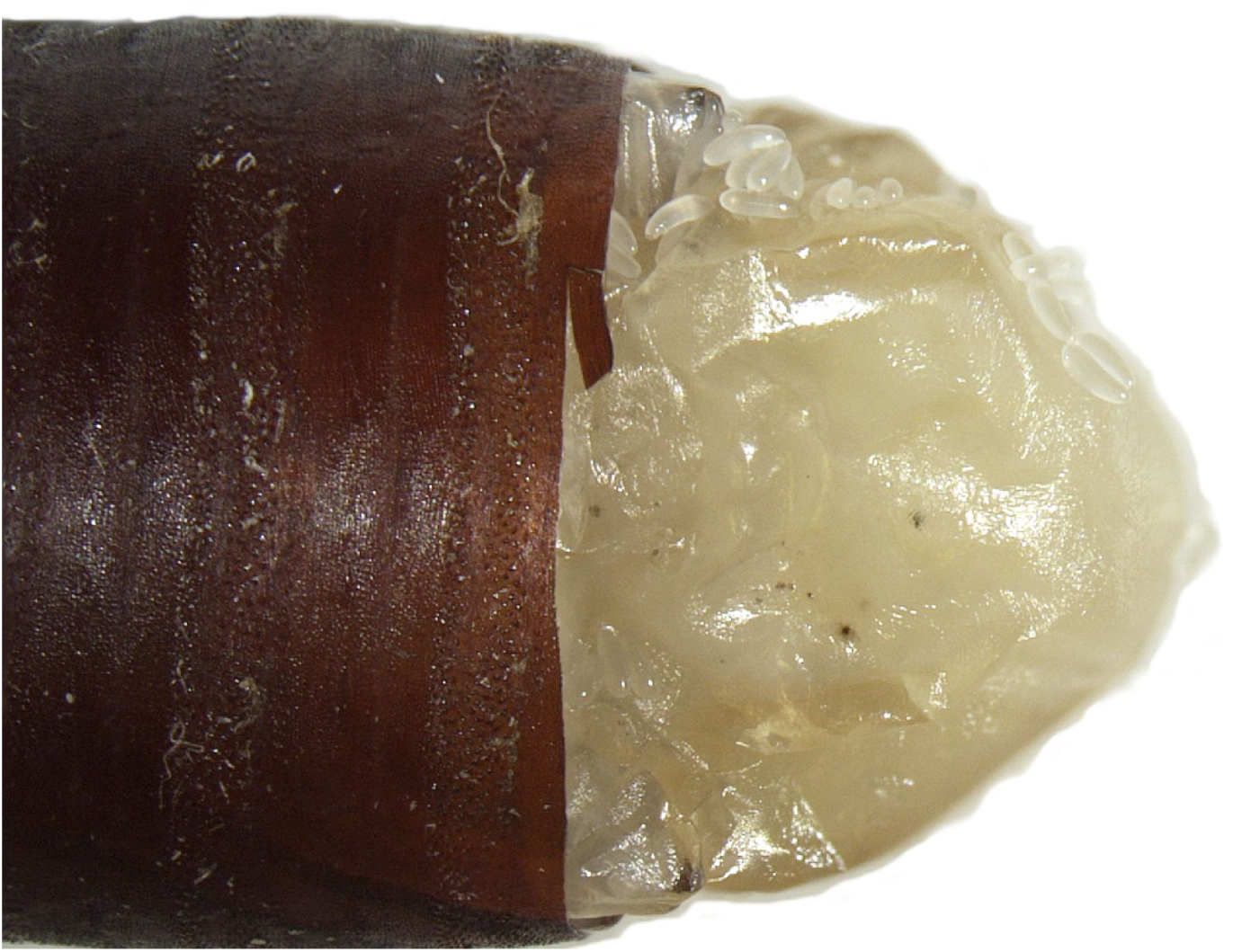
Parasitized *Sarcophaga bullata* pupae by *N. vitripennis.* The puparium has been peeled off at the posterior end using forceps, thereby exposing the *N. vitripennis* eggs.

**Figure 5.**
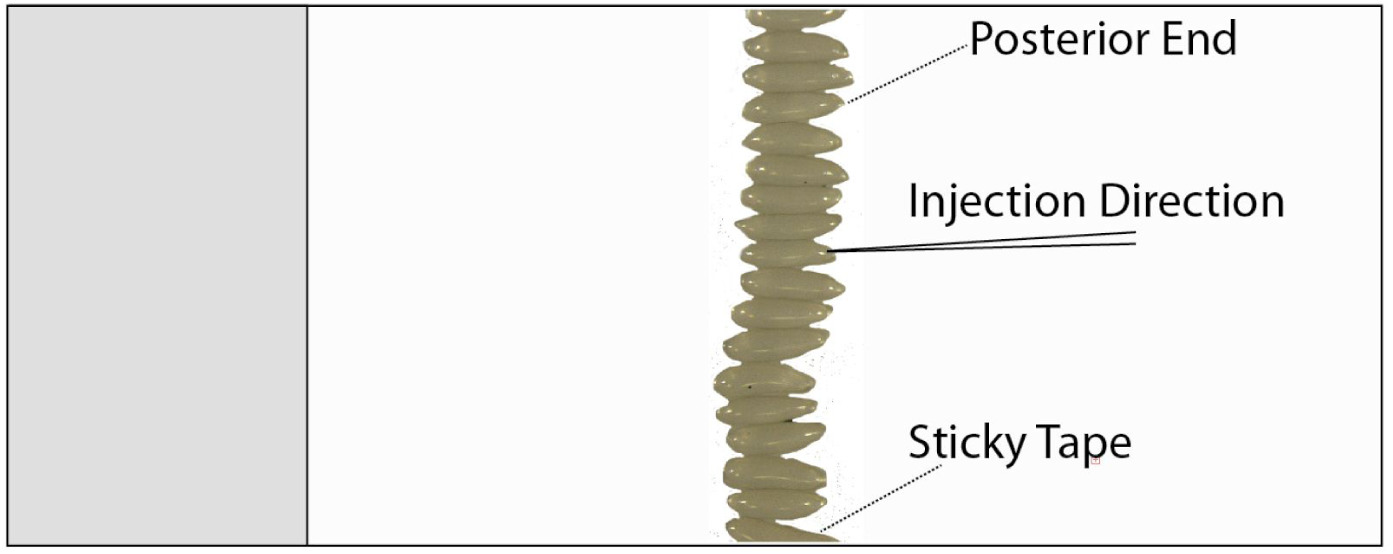
*N. vitripennis* embryos are lined up on a glass slide on a piece of sticky tape, and injected one-by-one into the posterior end.

**Figure 6.**
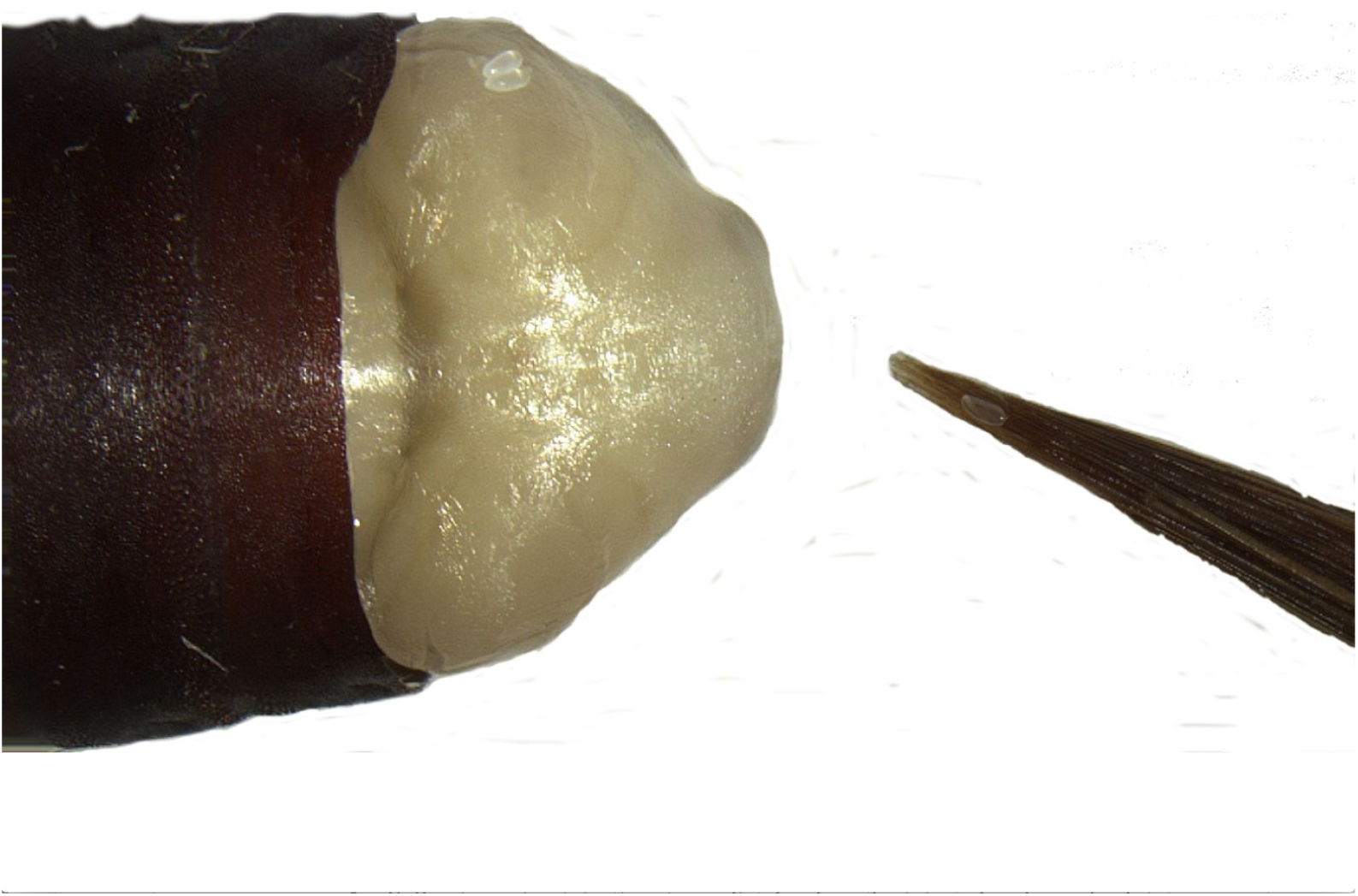
Injected *N. vitripennis* embryos are individually transferred to a pre-stung *Sarcophaga bullata* pupae using a wet fine-tip paintbrush. The puparium has been peeled off at the posterior end using forceps, to allow placement of the *N. vitripennis* eggs.

**Figure 7.**
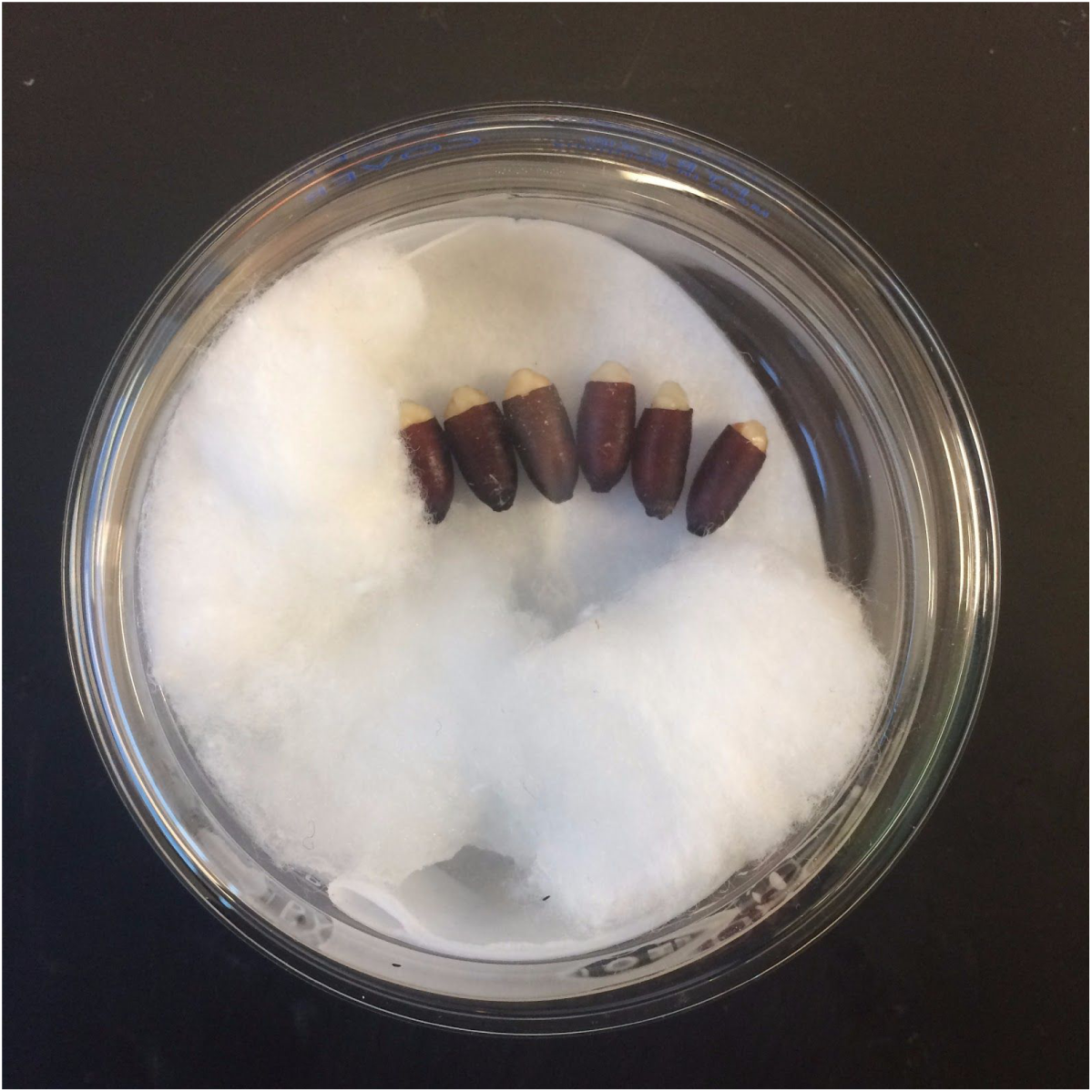
Incubation of *Sarcophaga bullata* hosts harboring the injected *N vitripennis* eggs in petri dish for roughly 2 weeks.

## References

1. Davis, R. B., Baldauf, S. L. & Mayhew, P. J. The origins of species richness in the Hymenoptera: insights from a family-level supertree. BMC Evol. Biol. 10, 109 (2010).

2. Whiting, A. R. The biology of the parasitic wasp Mormoniella vitripennis [= Nasonia brevicornis](Walker). Q. Rev. Biol. 333–406 (1967).

3. Ferree, P. M., Avery, A., Azpurua, J., Wilkes, T. & Werren, J. H. A bacterium targets maternally inherited centrosomes to kill males in Nasonia. Curr. Biol. 18, 1409–1414 (2008).

4. Hurst, G. D. & Jiggins, F. M. Male-killing bacteria in insects: mechanisms, incidence, and implications. Emerg. Infect. Dis. 6, 329–336 (2000).

5. Werren, J. H. & Stouthamer, R. PSR (paternal sex ratio) chromosomes: the ultimate selfish genetic elements. Genetica 117, 85–101 (2003).

6. Nur, U., Werren, J. H., Eickbush, D. G., Burke, W. D. & Eickbush, T. H. A ‘selfish’ B *chromosome that enhances its transmission by eliminating the paternal genome*. Science 240, 512–514 (1988).

7. Akbari, O. S., Antoshechkin, I., Hay, B. A. & Ferree, P. M. Transcriptome profiling of Nasonia vitripennis testis reveals novel transcripts expressed from the selfish B chromosome, paternal sex ratio. G3 3, 1597–1605 (2013).

8. Werren, J. H. et al. Functional and evolutionary insights from the genomes of three parasitoid Nasonia species. Science 327, 343–348 (2010).

9. Desjardins, C. A. et al. Fine-scale mapping of the Nasonia genome to chromosomes using a high-density genotyping microarray. G3 3, 205–215 (2013).

10. Ferree, P. M. et al. Identification of Genes Uniquely Expressed in the Germ-Line Tissues of the Jewel Wasp Nasonia vitripennis. G3 5, 2647–2653 (2015).

11. Sackton, T. B., Werren, J. H. & Clark, A. G. Characterizing the infection-induced transcriptome of Nasonia vitripennis reveals a preponderance of taxonomically-restricted immune genes. PLoS One 8, e83984 (2013).

12. Lynch, J. A. & Desplan, C. A method for parental RNA interference in the wasp Nasonia vitripennis. Nat. Protoc. 1, 486–494 (2006).

13. Werren, J. H., Loehlin, D. W. & Giebel, J. D. Larval RNAi in Nasonia (parasitoid wasp). Cold Spring Harb. Protoc. 2009, db.prot5311 (2009).

14. Beukeboom, L. & Desplan, C. Nasonia. Curr. Biol. 13, R860 (2003).

15. Shuker, D., Lynch, J. & Peire Morais, A. Moving from model to non-model organisms? Lessons from Nasonia wasps. Bioessays 25, 1247–1248 (2003).

16. Lynch, J. A. The expanding genetic toolbox of the wasp Nasonia vitripennis and its relatives. Genetics 199, 897–904 (2015).

17. Danneels, E. L., Rivers, D. B. & de Graaf, D. C. Venom proteins of the parasitoid wasp Nasonia vitripennis: recent discovery of an untapped pharmacopee. Toxins 2, 494–516 (2010).

18. de Graaf, D. C. et al. Insights into the venom composition of the ectoparasitoid wasp Nasonia vitripennis from bioinformatic and proteomic studies. Insect Mol. Biol. 19 Suppl 1, 11–26 (2010).

19. Verhulst, E. C., Beukeboom, L. W. & van de Zande, L. Maternal control of haplodiploid sex determination in the wasp Nasonia. Science 328, 620–623 (2010).

20. Bordenstein, S. R., O’Hara, F. P. & Werren, J. H. Wolbachia-induced incompatibility precedes other hybrid incompatibilities in Nasonia. Nature 409, 707–710 (2001).

21. Pultz, M. A. et al. A major role for zygotic hunchback in patterning the Nasonia embryo. Development 132, 3705–3715 (2005).

22. Lynch, J. A., Brent, A. E., Leaf, D. S., Pultz, M. A. & Desplan, C. Localized maternal orthodenticle patterns anterior and posterior in the long germ wasp Nasonia. Nature 439, 728–732 (2006).

23. Lynch, J. A., Olesnicky, E. C. & Desplan, C. Regulation and function of tailless in the long germ wasp Nasonia vitripennis. Dev. Genes Evol. 216, 493–498 (2006).

24. Werren, J. H. & Loehlin, D. W. The parasitoid wasp Nasonia: an emerging model system with haploid male genetics. Cold Spring Harb. Protoc. 2009, db.emo134 (2009).

25. Gaj, T., Gersbach, C. A. & Barbas, C. F., 3rd. ZFN, TALEN, and CRISPR/Cas-based methods for genome engineering. Trends Biotechnol. 31, 397–405 (2013).

26. Sethuraman, N. & O’Brochta, D. A. The Drosophila melanogaster cinnabar gene is a cell autonomous genetic marker in Aedes aegypti (Diptera: Culicidae). J. Med. Entomol. 42, 716–718 (2005).

27. Paton, D. R. & Sullivan, D. T. Mutagenesis at the cinnabar locus in Drosophila melanogaster. Biochem. Genet. 16, 855–865 (1978).

28. Cornel, A. J., Benedict, M. Q., Rafferty, C. S., Howells, A. J. & Collins, F. H. Transient expression of the Drosophila melanogaster cinnabar gene rescues eye color in the white eye (WE) strain of Aedes aegypti. Insect Biochem. Mol. Biol. 27, 993–997 (1997).

29. Labun, K., Montague, T. G., Gagnon, J. A., Thyme, S. B. & Valen, E. CHOPCHOP v2: a web tool for the next generation of CRISPR genome engineering. Nucleic Acids Res. 44, W272–6 (2016).

30. Xie, S., Shen, B., Zhang, C., Huang, X. & Zhang, Y. sgRNAcas9: a software package for designing CRISPR sgRNA and evaluating potential off-target cleavage sites. PLoS One 9, e100448 (2014).

31. Hsu, P. D. et al. DNA targeting specificity of RNA-guided Cas9 nucleases. Nat. Biotechnol. 31, 827–832 (2013).

32. Beukeboom, L. W. & van de Zande, L. Genetics of sex determination in the haplodiploid wasp Nasonia vitripennis (Hymenoptera: Chalcidoidea). J. Genet. 89, 333–339 (2010).

33. Zwier, M. V., Verhulst, E. C., Zwahlen, R. D., Beukeboom, L. W. & van de Zande, L. DNA methylation plays a crucial role during early Nasonia development. Insect Mol. Biol. 21, 129–138 (2012).

34. Özüak, O., Buchta, T., Roth, S. & Lynch, J. A. Dorsoventral polarity of the Nasonia embryo primarily relies on a BMP gradient formed without input from Toll. Curr. Biol. 24, 2393–2398 (2014).

35. Rosenberg, M. I., Brent, A. E., Payre, F. & Desplan, C. Dual mode of embryonic development is highlighted by expression and function of Nasonia pair-rule genes. Elife 3, e01440 (2014).

36. Bassett, A. R., Tibbit, C., Ponting, C. P. & Liu, J.-L. Highly efficient targeted mutagenesis of Drosophila with the CRISPR/Cas9 system. Cell Rep. 4, 220–228 (2013).

37. Bi, H.-L., Xu, J., Tan, A.-J. & Huang, Y.-P. CRISPR/Cas9-mediated targeted gene mutagenesis in Spodoptera litura. Insect Sci. 23, 469–477 (2016).

38. Doench, J. G. et al. Rational design of highly active sgRNAs for CRISPR-Cas9-mediated gene inactivation. Nat. Biotechnol. 32, 1262–1267 (2014).

39. Kohno, H., Suenami, S., Takeuchi, H., Sasaki, T. & Kubo, T. Production of Knockout Mutants by CRISPR/Cas9 in the European Honeybee, Apis mellifera L. Zoolog. Sci. 33, 505–512 (2016).

40. Gratz, S. J. et al. Highly specific and efficient CRISPR/Cas9-catalyzed homology-directed repair in Drosophila. Genetics 196, 961–971 (2014).

41. Kistler, K. E., Vosshall, L. B. & Matthews, B. J. Genome engineering with CRISPR-Cas9 in the mosquito Aedes aegypti. Cell Rep. 11, 51–60 (2015).

42. Hammond, A. et al. A CRISPR-Cas9 gene drive system targeting female reproduction in the malaria mosquito vector Anopheles gambiae. Nat. Biotechnol. 34, 78–83 (2016).

43. Gantz, V. M. et al. Highly efficient Cas9-mediated gene drive for population modification of the malaria vector mosquito Anopheles stephensi. Proc. Natl. Acad. Sci. U. S. A. 112, E6736–43 (2015).

## Supplemental References

1. Bassett, A. R., Tibbit, C., Ponting, C. P. & Liu, J.-L. Highly efficient targeted mutagenesis of Drosophila with the CRISPR/Cas9 system. Cell Rep. 4, 220–228 (2013).

2. Kistler, K. E., Vosshall, L. B. & Matthews, B. J. Genome engineering with CRISPR-Cas9 in the mosquito Aedes aegypti. Cell Rep. 11, 51–60 (2015).

3. Hwang, W. Y. et al. Efficient genome editing in zebrafish using a CRISPR-Cas system. Nat. Biotechnol. 31, 227–229 (2013).

4. Mali, P., Esvelt, K. M. & Church, G. M. Cas9 as a versatile tool for engineering biology. Nat. Methods 10, 957–963 (2013).

5. Ran, F. A. et al. Genome engineering using the CRISPR-Cas9 system. Nat. Protoc. 8, 2281–2308 (2013).

6. Wiedenheft, B., Sternberg, S. H. & Doudna, J. A. RNA-guided genetic silencing systems in bacteria and archaea. Nature 482, 331–338 (2012).

8. Ferree, P. M. et al. Identification of Genes Uniquely Expressed in the Germ-Line Tissues of the Jewel Wasp Nasonia vitripennis. G3 5, 2647–2653 (2015).

9. Labun, K., Montague, T. G., Gagnon, J. A., Thyme, S. B. & Valen, E. CHOPCHOP v2: a web tool for the next generation of CRISPR genome engineering. Nucleic Acids Res. 44, W272–6 (2016).

10. Xie, S., Shen, B., Zhang, C., Huang, X. & Zhang, Y. sgRNAcas9: a software package for designing CRISPR sgRNA and evaluating potential off-target cleavage sites. PLoS One 9, e100448 (2014).

11. Hsu, P. D. et al. DNA targeting specificity of RNA-guided Cas9 nucleases. Nat. Biotechnol. 31, 827–832 (2013).

13. Spradling, A. C. & Rubin, G. M. Transposition of cloned P elements into Drosophila germ line chromosomes. Science 218, 341–347 (1982).

